# On feature selection to disentangle cell type and state transcriptional programs

**DOI:** 10.1101/2024.11.29.626057

**Authors:** Jiayi Wang, Helena L. Crowell, Mark D. Robinson

## Abstract

Single-cell omics approaches profile molecular constituents of individual cells. Replicated multi-condition experiments in particular aim at studying how the molecular makeup and composition of cell subpopulations changes at the sample-level. Two main approaches have been proposed for these tasks: firstly, cluster-based methods that group cells into (non-overlapping) subpopulations based on their molecular profiles and, secondly, cluster-free but neighborhood-based methods that identify (overlapping) groups of cells in consideration of cross-condition changes. In either approach, discrete cell groups are subjected to differential testing across conditions; and, a low-dimensional cell embedding, which is in turn derived from a subset of selected features, is required to delineate subpopulations or neighborhoods.

We hypothesized that decoupling differences in cell type (i.e., between subpopulations) and cell state (i.e., between conditions) for feature selection would yield an embedding space that captures different aspects of cellular heterogeneity. And, that type-not-state embeddings would arrive at differential testing results that are more comparable between cluster- and neighborhood-based differential testing approaches. Our study leverages a simulation framework with competing type and state effects, as well as an experimental dataset, to evaluate a set of feature scoring and selection strategies, and to compare results from downstream differential analyses.

## Introduction

Single-cell RNA sequencing (scRNA-seq) and other related single-cell or multi-omics approaches now represent a robust suite of technologies for unraveling molecular phenotypes at cell-level resolution^[1]^. A now-common experimental design involves the comparison of molecular profiles from multi-subpopulation samples across conditions (e.g., across disease states, or before and after perturbation). If it is straight-forward to assign cells to (discrete) subpopulations (i.e., cell types or subtypes; see Discussion below), some fundamental discovery analyses include differential abundance analysis (DAA) to identify a change in subpopulation composition between conditions, or differential state analysis (DSA) to investigate changes in gene expression for a specific subpopulation across conditions; for the latter, pseudobulking the scRNA-seq data and leveraging classical differential expression (DE) frameworks^[2,3,4]^ in a way that accounts for sample-to-sample variability has been argued to result in good performance in terms of sensitivity and error control^[5,6,7]^.

Returning to the thorny issue of assigning cells to cell types, one can think of a cell type as a relatively permanent aspect (i.e., a set of stably expressed genes), whereas the cell state represents a more transient phenotype (i.e., another set of genes expressed temporarily)^[8,9]^. In a recent review that proposed to separate cell types from cell states, Xia & Yanai^[10]^ “view a cell state as a secondary module operating in addition to the general cell type regulatory program”. This statement is in tune with standard single-cell analysis pipelines, which generate low-dimensional embeddings from a set of highly variable genes^[11,12,13]^; such embeddings are thus influenced by transcriptional programs characteristic to both cell type and cell state, although the dominant signal is usually assumed to stem from the type program^[10]^.

Discretization into cell subpopulations itself has advantages and disadvantages. On the one hand, discretization makes for relatively clear interpretation, assuming the subpopulations are decorated with markers. However, an immediate tension is the appropriate resolution of the subpopulations, which is often subjectively specified. Furthermore, DE at broad resolution could simply reflect compositional changes of cell subtypes, whereas a too finely-grained resolution could lead to limits on statistical power. A few possibilities have now emerged to address the resolution problem. Our own work, treeclimbR, proposed to over-cluster into a higher resolution and, using a tree of relationships between clusters, aggregate the differential statistics in a data-driven fashion^[14]^; challenges remain to specify what set of features to over-cluster on and how to build a tree to aggregate results from. Related work looks at DE via dimension reduction (e.g., eSVD-DE ^[15]^) or the change in percentage of cells expressed^[16]^, but the starting point is still a set of clusters or annotated cell types. Meanwhile, a handful of “cluster-free” approaches have now been proposed for DSA. For example, miloDE ^[17]^ extends miloR’s strategy (for DAA^[18]^) to compute DE at local neighborhoods instead of clusters; neighborhoods are retained according to a minimum size to increase statistical power, while standard DE models can be used to correct for covariates. MELD ^[19]^ and Cacoa^[20]^ compute DE at the single-cell level (or rather in a neighborhood around each cell), but are unable to adjust for covariates. However, all these strategies typically use a low-dimensional embedding that mixes cell type and cell state effects. LEMUR^[21]^ presents a unified approach that uses a common low-dimensional latent space from all cells and all conditions to select neighborhoods (for each gene) that exhibit DE between conditions.

In this work, we investigated feature selection as a means to separate cell type and cell state programs; this strategy mimics somewhat the case of targeted single-cell omics (e.g., flow and mass cytometry), where markers are typically selected in advance to represent cell types or states^[22]^. We hypothesized that focusing gene selection on features predominantly contributing to cell type regulatory programs would yield embeddings less affected by cell state effects; and by doing this, results from downstream DSA would be more comparable between cluster-based and cluster-free approaches.

## Results

### Simulation setup

To investigate the entanglement of type and state effects in scRNA-seq data, we first relied on synthetic data to explore feature scoring and selection strategies, data properties post processing, and downstream DSA results (Fig. 1a). Specifically, we used Splatter ^[23]^ to simulate data across a grid of two competing simulation parameters, *t* and *s*, that control the similarity between cells according to clusters (types) and conditions (states), respectively (simplified schematic in Fig. 1b, full grid in Fig. S1). These parameters control the location parameter of log-normal distributions from which pair-wise fold-changes between clusters (*t*) and conditions (*s*) are sampled (Fig. 1c, Fig. S2). Because genes are selected randomly for each effect, they may exhibit changes in type or state, or both. For every pair of parameters, data for three clusters and three replicates from two conditions were simulated; each simulation was processed in a standardized manner, i.e., removal of low-quality genes and cells, library-size normalization, highly variable feature selection, principal component analysis (PCA), low- and high-resolution clustering.

**Fig. 1:**
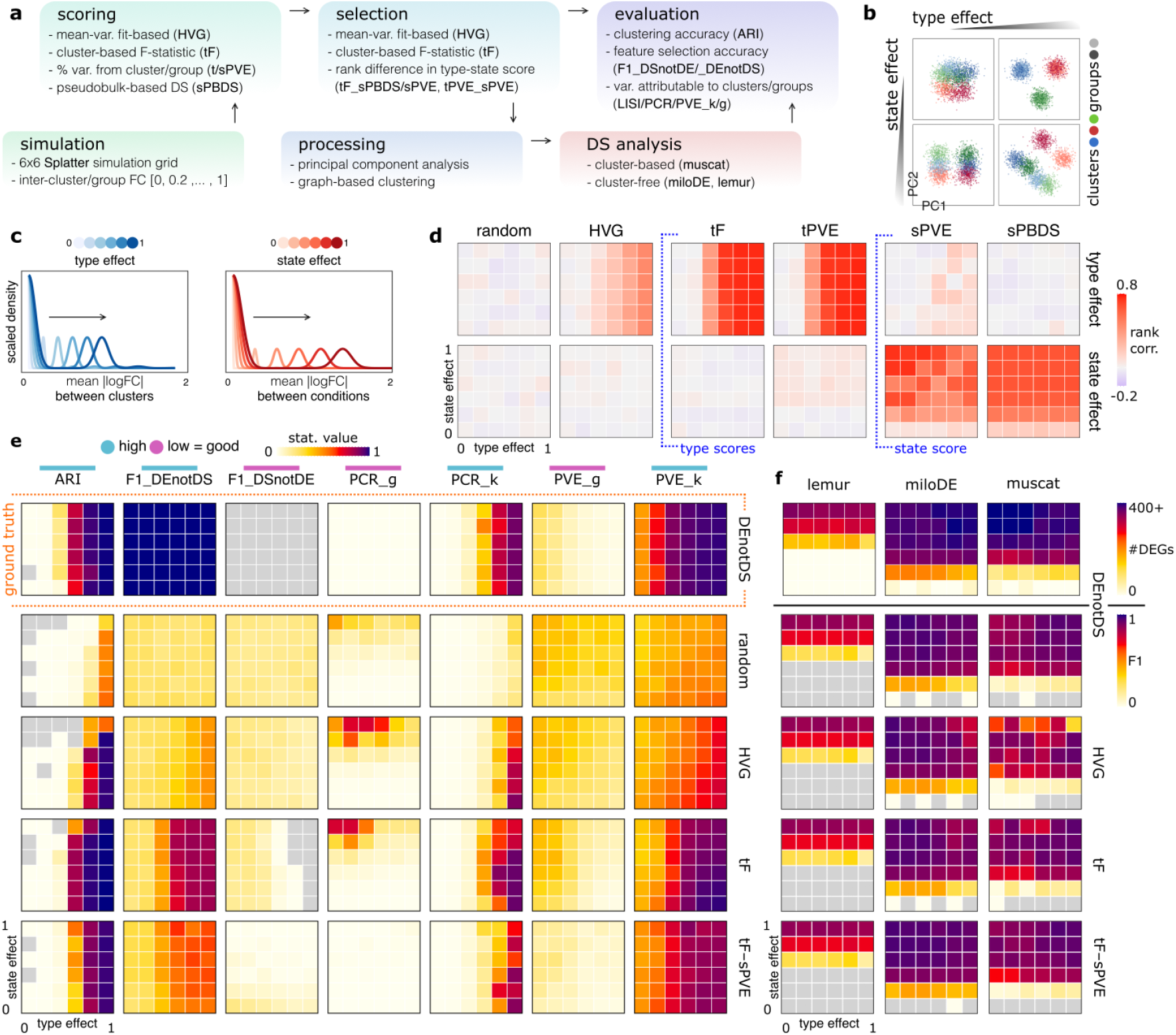
Simulation study design and results. (**a**) Simulation study workflow; methods and metrics employed at each step are highlighted in bold. (**b**) Scatter plots of first and second principal components (PCs) for exemplary simulations with varying type and state effects, i.e., degree of inter-cluster and -group expression dissimilarity. (**c**) Schematic of gene categorizations in a multi-sample and -group single-cell data setting. (**d**) Rank correlation between gene-wise simulated fold changes (FCs) between clusters (top row) or groups (bottom row) and gene-wise feature scores (columns) calculated for the simulated data. (**e**) Evaluation statistics computed against the ground truth (columns) across feature selections (rows) for each simulation; whether a high or low score is desirable is indicated by the color bar avoid the label. (**f**) Number of genes called differential and DS-based F1 scores (adjusted p-value < 0.05 in any comparison) across feature selections (rows) and DS analysis methods (columns). Each cell in a heatmap in **d, e** and **f** corresponds to a level of simulated type and state effects.

### Feature scoring

To quantify variation in gene expression attributable to types and states, respectively, we implemented a number of feature-wise scores (Table S1). These scores aim to measure either “typeness” (i.e., expression limited to one or a small number of subpopulations, and stable expression across conditions) or “stateness” (i.e., subpopulation-specific expression changes across conditions). Such scores can then be combined to select for type features and select against state features.

For type scores, we considered moderated F-statistics (tF) that score whether gene expression deviates across clusters, and the percent variance explained by cluster assignments (tPVE). For state scores, we considered the PVE by condition (sPVE) as well as DSA adjusted p-values from pseudobulks (sPBDS; e.g., Crowell *et al*.^[5]^). Lastly, we used a standard gene selection: highly variable genes (HVG)^[24]^. Notably, tF and PVE-based scores use high-resolution, while sPBDS is calculated on low-resolution cluster assignments. For reference, a random score was also included.

To judge how well these feature scores capture typeness and stateness, we correlated them against ground truth measures from the simulation (Fig. 1d). Here, “true” typeness is an average of the cluster-wise fold-changes (a reflection of changes in the mean between clusters), and “true” stateness is an average of condition-specific fold-changes (a reflection of changes in the mean between conditions within clusters); see Fig. 1c and Eq. S1. Correlations of true typeness with HVG and type scores (tF, tPVE) all increase with *t*, but are mostly invariant to changes in *s*. Vice versa, correlations of true stateness with state scores (sPVE, sPBDS) increased with *s* and are much less affected by *t*. However, tPVE and sPVE are not entirely robust to changes in *s* and *t*, respectively. Random scores do not correlate with either type of effect at any magnitude.

### Feature selection

In addition to random, HVG and tF score-based selection, we implemented three strategies to select for type-specific features that explicitly account for changes in state (Table S2). For example, tF-sPBDS selects features based on difference in ranks of tF and sPBDS scores, thereby prioritizing genes with high type and low state scores; tF-sPVE and tPVE-sPVE use analogous strategies. Finally, we processed simulations independently for each feature selection strategy (PCA on a fixed number of selected features, graph-based clustering, and DSA).

### Evaluation statistics

To evaluate feature selection, we computed metrics that assess retrieval of the simulated clusters or of type/state genes, as well as the global data structure (see Table S3 for verbose descriptions); for example, we used: adjusted rand index (ARI) between simulated and clusters retrieved from data processing after gene selection; F1 scores of selected features according to various definitions of ground truth (e.g., DEnotDS denotes the set of genes that are DE between clusters but not across conditions; further definitions are given in Table S3); the local inverse Simpson index^[25]^ considering *g*/*k* as variables (LISI_g, LISI_k) to quantify mixing of cells from different labels; the explained variance-weighted sum of coefficients of determination from principal component regression using *g*/*k* as predictor (PCR_g, PCR_k) to quantify the amount of variability attributable to each; and, the PVE^[26]^ by *g*/*k* (PVE_g, PVE_k). Here, LISI and PCR are computed on PCs of selected features, while PVE is based on their expression values.

First, we evaluated feature selections that are based on simulated effects between types and states; specifically: DE/DS (type/state effect is present), DEgtDS/DSgtDE (on average, type/state effect exceeds state/type effect), and DEnotDS/DSnotDE (type/state effect, but *no* state/type effects) (Table S2). Overall, DEnotDS performed best across all evaluation criteria; specifically, cluster-based statistics (ARI, PCR_k, PVE_k, LISI_k) improved with the type effect (*t*) but are invariant to state changes (*s*), while condition-based statistics (PCR_g, PVE_g, LISI_g) are at an optimum for most simulations (Fig. S4a, Fig. S5); hence, we selected the top-*n* ranked features for all strategies, where *n* corresponds to the number of DEnotDS genes in a given simulation.

Random and HVG selections were worst overall and most sensitive to state changes (*s*); for example, F1_DEnotDS is ≈ 0.5 for most simulations, PVE_k decreases and PCR_g/PVE_g increase with *s* (Fig. 1e, Fig. S4). Owing to its exclusive consideration of type effects, tF performs well in cluster-based metrics (ARI, PCR_k, PVE_k), but it shows poorer performance in group-based metrics (PCR_g, PVE_g). Methods considering both type and state effects (tF-sPBDS, tF-sPVE and tPVE-sPVE) gave results most similar to those obtained for ground truth-based selection (DEnotDS). When *t* is sufficiently large, these selections were able to resolve clusters based on the first couple of PCs while keeping cells from different conditions together (Fig. S6). In contrast, HVG selection separated cells by condition in many cases.

### Differential discovery

Next, we subjected data processed through various feature selections to DSA using muscat (i.e., at pseudobulk-level) and two recent cluster-free approaches, miloDE ^[17]^ and lemur ^[21]^; notably, lemur is unaffected by gene selection and instead learns a latent space representation of cell type and state. In any DSA, each gene undergoes testing across a set of ‘instances’. For cluster-based methods, these are *disjoint* subpopulations; for cluster-free methods, they may *overlap* and correspond to groups of cells with similar state changes. Consequently, it is challenging to directly compare their results. Here, we extracted the minimum adjusted p-values (denoted p^∗^) for each gene across clusters (muscat) or neighborhoods (lemur, miloDE); genes with p^∗^ < 0.05 in any comparison were considered ‘hits’. miloDE gave more accurate results than muscat, and was largely robust to changes in simulation parameters; lemur was notably less sensitive and unable to identify DS genes when *s* < 0.6 (short list of selections in Fig. 1f; full list in Fig. S7, Fig. S8). Correlating p^∗^ between DSA methods yields reasonable concordance between muscat and miloDE, but not lemur against other methods (Fig. S9).

### Experimental data

To see how these findings manifest in a real dataset, we analyzed the scRNA-seq data from Kang *et al*.^[27]^, which profiles PBMCs from Lupus patients before and after a 6-hour IFN-*β* treatment. We selected the top 2,000 genes for each gene selection strategy and processed the data in a standard way (PCA, graph-based clustering, and DSA). Low-dimensional projections of cells using random, HVG and tF show clear separation of cells between conditions, even when they belong to the same cell type (Fig. 2a, Fig. S10), thus highlighting the mix of type and state programs within the embedding. However, a type-not-state feature selection strategy (tF-sPBDS) retained primarily the cell type structure. This is a reflection of PC2 capturing cell state rather than cell type differences for random, HVG, and tF selection, but not for tF-sPBDS (see Fig. 2b; full list in Fig. S11). Quantitative evaluations (based on ARI, PCR and PVE, which do not require a feature-level ground truth) showed that type-not-state gene selection better recovered the annotated cell types as clusters (ARI, LISI_g, LISI_k; see Fig. 2c). Additionally, these selections attribute more variation to cell type than to condition (PCR_g, PCR_k and PVE_g, PVE_k). Notably, many of the evaluation metrics are affected by the number of features selected (Fig. S12) and factors such as the number of PCs and clustering resolution will affect the number of recovered clusters and the performance of different selections.

**Fig. 2:**
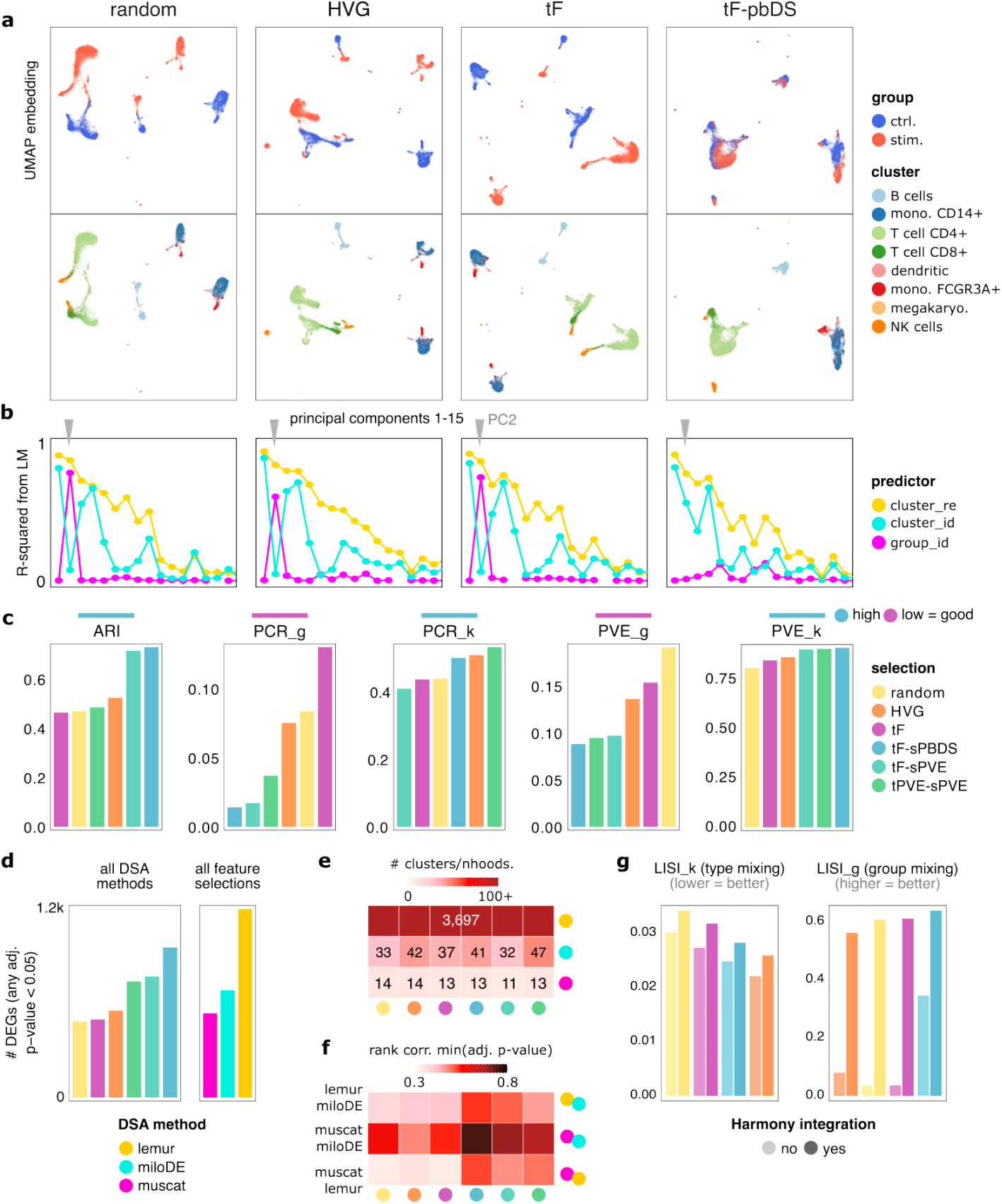
Experimental data results. (**a**) UMAP embeddings of the Kang *et al*^.[27]^ dataset based on PCs computed from different feature selections. Points (= cells) are colored by experimental condition (top) and cell type (bottom), respectively. (**b**) Coefficients of determination (R-squared) from fitting a linear model (LM) against principal components (PCs) with clusters (cluster_re), annotated subpopulations from Kang *et al*.^[27]^ (cluster_id), and experimental condition (group_id) as predictor, respectively. (**c**) Comparison of evaluation statistics across feature selections. (**d**) Number of genes with an adjusted *p*-value < 0.05 in any comparison (cluster or neighborhood) called by all DSA methods (stratified by selection), and called for any feature selection method or random selection only (stratified by DSA method). (**e**) Number of groups identified for testing across selections and DSA methods (neighborhoods for lemur and miloDE, clusters for muscat). (**f**) Spearman rank correlation between smallest adjusted *p*-values from different pairs of DSA methods (rows), stratified by feature selections (columns). (**g**) Comparison of PCA-based LISI_k and LISI_g statistics across feature selections, with and without using Harmony to correct for batch effects across samples.

Lacking ground truth, we compared the number and concordance of genes deemed differential by DSA methods for a given feature selection (Fig. 2d; c.f. Fig. S13). tF-based selection and HVGs resulted in < 500 common DEGs, whereas selections based on type-not-state scores resulted in *>* 500 − 1000 common DEGs (i.e., DSA methods have higher concordance); random selection ranked similar to HVGs. miloDE returned the most DEGs for *any* selection (followed by lemur, then muscat), i.e., the methods results are most robust to the features selected for. DSA methods differed greatly in the number of tested instances. Being cluster-based, muscat was closest to the ‘true’ number of subpopulations (8); miloDE detected 10s of neighborhoods, and lemur considered a unique neighborhood for most genes (Fig. 2e). The correspondence of these instances to cell ‘types’, however, will arguably depend on the chosen resolution (e.g., subpopulation definitions are rather broad in these data). Similar to simulated data, the DSA results of the Kang dataset followed our hypothesis that cell embeddings based primarily on type (and not state) programs lead to more concordant DSA results. This can be observed through the higher correlation between different DSA methods, especially muscat and miloDE, when using feature selections that combine type and state scores (Fig. 2f).

Lastly, we quantified the mixing of clusters and groups across feature selections, as well as before and after Harmony^[25]^ integration to correct for batch effects between samples (Fig. 2g). While HVGs give the best separation of clusters, they also separate groups, which is suboptimal. Integration markedly improves group mixing but also increases mixing of clusters. Combining integration with type-not-state selection gives the best result overall. Thus, feature selection and integration represent complementary steps.

## Discussion

There is arguably an interplay between not only the type and state transcriptional programs governing cells, but also between the various types of differential analysis that are applied. While the readily-available DAA and DSA tools can in principle be considered orthogonal, they are often linked by how DE drives subpopulation definition^[17]^. As a toy example, consider a substance that strongly increases the expression of a gene *γ* in all treated (but not control) cells. If *γ* were used for clustering (or neighborhood generation), *γ*^−^ and *γ*^+^ subpopulations should be obtained for each cell type; and, DAA analysis would detect *γ*^+^ and *γ*^−^ populations as differentially abundant. In contrast, if *γ* were *not* contributing to the subpopulation definition, *γ*^+^ and *γ*^−^ cells should be present in every cluster (whose markers are expressed independently from *γ*), and a DSA analysis should detect *γ* as higher expressed in treated cells (i.e., a differential state) for each cell type.

Ultimately, both DAA and DSA could detect a defined biological effect (e.g., inflammation), regardless of how embeddings (and therefore subpopulations) are defined. However, we highlighted here that a key factor influencing interpretation, regardless of whether cluster- or neighborhood-based approaches are used, is the set of features contributing to the input embeddings. By separating type and state patterns for feature selection, we can not only simplify interpretation, but also derive results where cluster- and neighborhood-based approaches are more concordant. This separation also provides a clearer input embedding for integration and could serve as a complementary component to existing integration methods. In place of highly variable genes, we suggest to isolate variation that is indicative of cell type programs to start downstream analyses with “clean” (type-not-state) embeddings.

Of course, further experience needs to be gained across various datasets and designs to understand how to best fine tune such feature selection, especially across the spectrum of discrete to continuous cell types^[28]^. Furthermore, the effect of type-not-state feature selection on integration across datasets needs to be investigated and in particular, when mapping datasets to reference atlases spanning multiple disease states.

The type/state models we tested are highly simplified: type features are those that are highly expressed in a small number of subpopulations across samples; state features are differentially expressed across conditions within subpopulations. The conundrum here is that we need cell type definitions in order to quantify cell state variation. Thus, a concrete next step would be to investigate whether iterative selection can reveal more stable features and thus more refined embeddings and subpopulation definitions.

In summary, whether cluster- or neighborhood-based approaches are used for differential discovery in the multi-sample multi-condition multi-subpopulation single cell datasets, a key consideration for interpretable data analysis is feature selection. We propose that isolating type-not-state variation directly into cell embeddings and subpopulation definitions will be a promising approach to unravel complex but interpretable subpopulation-specific effects^[29]^.

## Data availability

The original Kang *et al*.^[27]^ scRNA-seq data is deposited in the Gene Expression Omnibus (accession GSE96583). A corresponding SingleCellExperiment object is available in R through Bioconductor’s ExperimentHub^[30]^ (identifier Kang18 _8vs8).

## Code availability

All analyses were run in R v4.4.1^[31]^ with Bioconductor v3.20^[32]^. The computational workflow was implemented using Snakemake v7.26.0^[33]^, with Python v3.11.3. Snakemake workflow and underlying R code are accessible at https://github.com/HelenaLC/type-state, as well as Zenodo (https://doi.org/10.5281/zenodo.14191289).

## Acknowledgements

The authors thank members of the Robinson Lab at the University of Zurich for valuable feedback on methodology and exposition, as well as Dr. Constantin Ahlmann-Eltze, author of lemur, for support regarding its usage. This work was supported by Swiss National Science Foundation (SNSF) project grant 310030_204869 to MDR. HLC acknowledges support by SNSF grant number 222136. MDR acknowledges support from the University Research Priority Program Evolution in Action at the University of Zurich.

## Authors’ contributions

HLC conceived the study design and the computational workflow. JW and HLC implemented methods, with assistance from MDR regarding methodology. All authors contributed to drafting the paper, and have read and approved its final version.

## Methods

### Feature scoring

#### Highly variable genes (HVG)

We used modelGeneVar (scran^[24]^) to model the variance of gene expression profiles, which returns estimates for technical and biological variance components based on a fitted mean-variance trend; the latter were used as HVG scores.

#### Cluster-based F-statistic (tF)

The F-statistic of a gene is calculated by the ratio of between-cluster variance to within-cluster variance. Rather than implementing the traditional F-statistics, we fitted a linear model (LM) using limma on sample-level pseudobulks, and empirical Bayes moderated F-statistics to summarize the gene expression variation between clusters, while accounting for sample-level effects. Here, high-resolution cluster assignments were used as input (i.e., from Louvain clustering with resolution parameter 2).

#### Percent variance explained by type/state (tPVE/sPVE)

To estimate the fraction of variance attributable to sample, group and cluster variables, respectively, we used fitExtractVarPartModel (variancePartition^[26]^) to fit a linear mixed model (LMM) according to *Y* ~ (1|*s*) + (1|*g*) + (1|*k*), where *Y* denotes gene expression (log-library size normalized counts), and *s/g/k* denote sample/group/cluster assignments. Again, high-resolution cluster assignments were used to fit the model. The PVE by clusters and groups are used as type scores (tPVE) and state scores (sPVE), respectively.

### Pseudobulk-based DSA (sPBDS)

Pseudobulk-based DSA was performed as proposed in previous work of ours^[5]^. Briefly, for every (low-resolution) cluster, sample-level pseudobulks (sum of counts) were subjected to cross-group comparison using edgeR^[2]^ with glmQLFit and glmQLFTest ^[34]^. Because genes can undergo multiple comparisons (when there are multiple clusters), we used −*log*(p^∗^) as sPBDS scores, where p^∗^ denotes the smallest adjusted p-value across clusters.

### Feature selection

We ranked genes based on different scores and selected the top-*n* ranked genes for reprocessing, where *n* is the number of the ground truth *type* genes (DEnotDS; c.f. Table S2). For individual scores, such as tF, the rank is defined as the decreasing order of score values. For combined scores, such as tF-sPVE, the rank is defined by the decreasing order of the rank difference between the type and state score, i.e., rank (tF)-rank (sPVE).

### Evaluation statistics

#### Adjusted rand index (ARI)

The ARI was computed between simulated (true) and reprocessed (predicted) cluster labels. Here, low values (close to 0) indicate random agreement, while high values (close to 1) indicate perfect concordance between true and predicted labels; for visualization, negative values were set to N/A.

#### F1_DEnotDS/F1_DSnotDE

The F1 score is a classification accuracy metric, computed as the harmonic mean of precision and recall, i.e., 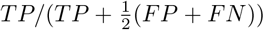, where TP, FP, and FN are the number of true positives, false positives, and false negatives, respectively. Here, F1 scores were computed between features selected by a given method, and features defined as “true” (e.g., *type* genes according to DEnotDS or *state* genes according to DSnotDE).

#### Local inverse Simpson’s index (LISI)

The LISI^[25]^ assesses the extent of mixing among clusters of cells across clusters (types) or conditions (states), denoted LISI_g and LISI_k. LISI assigns a cell-level diversity score, indicating the effective number of different cell types (*LISI k*) or conditions (*LISI g*) represented in the local neighborhood of each cell. For example, a well-mixed scenario is indicated by a LISI score close to 2 for a binary categorical variable, while a well-separated case is indicated by a LISI score close to 1 regardless of the number of categorical variables.

For visualization, LISI-based scores *L* are scaled between 0 and 1 via (*L* − 1)*/*(*n* − 1), where *n* is the number of categorical variables, i.e., the number of clusters/conditions for LISI_k/LISI_g.

#### Percent variance explained by group/cluster (PVE_g/PVE_k)

fitExtractVarPartModel (variancePartition^[26]^) was again used for evaluating the fraction of gene expression variance attributable to a gene’s cluster (type) and group (state), denoted PVE k/PVE g, by fitting a LMM according to *Y* ~ (1|*g*) + (1|*k*), where *Y* denotes log-library size normalized counts, and *k/g* denotes cluster/group assignment. Estimates were then re-scaled such that they sum to 1 disregarding residuals.

#### Principal component regression against groups/clusters (PCR g/PCR k)

We regressed each principal component *PC* = {*pc*_1_, …, *pc*_*N*_} against a cell-level variable *j* of interest; here, group and cluster assignments, respectively. The resulting coefficients of determination R^2^ were then weighted by the variance in gene expression *Y* (log-library size normalized counts) attributable to the corresponding PC, summed across PCs, and scaled by the overall variance, i.e.:

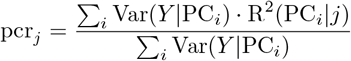

### Differential state analyses

lemur ^[21]^ and miloDE ^[17]^ present neighborhood-based approaches to DSA, while muscat ^[5]^ relies on clustering of cells from which to derive sample-level pseudobulks that are subjected to differential testing; both types of methods rely on a prior low-dimensional embedding of cells. In order to compare these methods’ results despite their methodological differences, we passed the same set of PCs to both miloDE and muscat ; these, in turn, depend on the upstream feature selection strategy. For miloDE, PCs were used as the embedding for graph construction; for muscat, pseudobulks subjected to differential testing with edgeR were based on clusters obtain from the same PCs. By contrast, the expression-level data was passed to lemur and corrected across groups via align_harmony ; latent embeddings (representing cell type and state heterogeneity) and cellular neighborhoods (potentially unique for every gene) are identified independently from previously selected features. All methods were run using default parameters.

### Experimental data

The Kang dataset is available through Bioconductor’s ExperimentHub with the accession ID EH2259 and contains 35,635 genes and 29,065 cells. We removed unassigned and multiplet cells according to authors’ annotation and filtered lowly expressed genes, i.e., genes detected in fewer than 20 cells. After filtering, we obtained 4,180 genes and 12,516 cells with 5,836 and 6,680 cells before and after treatment, respectively, annotated into eight subpopulations. Next, we preprocessed the data and scored each feature using the same strategy as for simulated data (see Supplementary Methods).

