## Supplementary Information for "On feature selection to disentangle cell type and state transcriptional programs"

### 1 Supplementary Tables

#### 1.1 Feature scores

| Scoring | Description | Interpretation |
|---|---|---|
| tF | Differential gene expression between cell types/clusters* (blocking on samples) using a linear model fit to cell-level log-normalized counts using limma, with summarization of 'typeness' using moderated F-statistics | type score |
| sPBDS | Pseudobulk-based DSA with edgeR, with summarization of 'stateness' using the minimum $p$-value across clusters* | state score |
| sPVE | Fraction of variance attributable to conditions (cell states) given by a linear mixture model on log-normalized counts | state score |
| tPVE | Fraction of variance attributable to clusters (cell types) given by a linear mixed model | type score |
| HVG | scran's modelGeneVar() bio estimates (biological component of variance) | standard gene selection score |
| random | $\sim \mathcal{U}_{[0,1]}$ | - |

**Table S1: Feature scoring methods.** Each gene receives one (aggregated) score from the list of scoring methods to quantify its typeness and stateness. *tF and PVE-based scoring methods reply on HVG-based high-resolution clustering, while sPBDS replies on low-resolution clustering (see Section 3.1).

For simulated data, a differential expression (DE) factor is assigned to each gene and each cluster or condition. Here, $i \in \{1, ..., N\}$ represents genes, $x \in \{1, ..., X\}$ indicates a group of cells, and $\delta_{ix}$ represents the DE factor induced by splatter on the $i$-th gene in the $x$-th group (in comparison to the baseline expression level). The simulated type or state effect (i.e. true typeness and stateness) for gene $i$ is quantified by the average log fold change:

$$logFC_i = \frac{1}{\binom{X}{2}} \sum_{x \neq x'} |\log(\frac{\delta_{ix'}}{\delta_{ix}})| \quad (1)$$

For type effect ($logFC_{type}$), $x \in \{1, 2, 3\}$ represents cell types; for state effect, ($logFC_{state}$), $x \in \{1, 2\}$ represents conditions. The true type and state effect in logFCs derived from the splatter simulation parameters are used to define selection strategies for establishing the ground truth across different expectations in Table S2.

#### 1.2 Feature selection

| Selection | Description | Interpretation |
|---|---|---|
| DEnotDS* | $logFC_{type} \neq 0$ and $logFC_{state} = 0$ | genes with type effects but without state effects |
| DEgtDS | $logFC_{type} > logFC_{state}$ | type effect greater than state effect |
| DE | $logFC_{type} \neq 0$ | type effect present |
| DSnotDE | $logFC_{state} \neq 0$ and $logFC_{type} = 0$ | state effect but no type effect |
| DSgtDE | $logFC_{state} > logFC_{type}$ | state effect greater than type effect |
| DS† | $logFC_{state} \neq 0$ | state effect present |
| random | based on score $\sim \mathcal{U}_{[0,1]}$ | – |
| HVG | based on biological component of variance | standard feature selection |
| tF | based on tF score only | high type score but ignores state score |
| tF-sPBDS | difference in ranks of tF and sPBDS | high type score but low state score |
| tF-sPVE | difference in ranks of tF and sPVE | same as above |
| tPVE-sPVE | difference in ranks of tPVE and sPVE | same as above |

**Table S2: Feature selection strategies** The first six rows (DEnotDS to DS) represent selection strategies based on simulation parameters for defining ground truth across different expectations. The remaining rows represent selection strategies based on scores calculated from the gene expression data. HVG is the standard feature selection used in scRNA-seq data; tF corresponds to DE; tF-sPBDS, tF-sPVE, and tPVE-sPVE correspond to DEnotDS or DEgtDS. (* = ground truth for feature selection; † = ground truth for DSA).

#### 1.3 Evaluation statistics

| Statistic | Description | Interpretation |
|---|---|---|
| ARI | (cluster-based) adjusted rand index | similarity between ground truth and reprocessed data cluster assignments (higher = better) |
| F1_DEnotDS | Using DEnotDS as gene-based ground truth; harmonic mean of precision and recall | accuracy in ground truth *type* gene retrieval (higher = better) |
| F1_DSnotDE | same as above but using DSnotDE as gene-based ground truth | accuracy in ground truth *state* gene retrieval (lower = better) |
| LISI_g/k | Inverse Simpson's Index to compute diversity of each cell's local neighborhood | mixing of groups/clusters (higher/lower = better) |
| PCR_g/k | sum over $R^2$ values from PC regression on groups/clusters weighted by each PCs PVE | (PC-based) variance attributable to groups/clusters (higher/lower = better) |
| PVE_g/k | (gene expression-based) variance attributable to groups/clusters | higher/lower = better |

**Table S3: Evaluation statistics.** $g$-based statistics are associated with groups (conditions/cell states), where a good selection is expected to result in low values; $k$-based statistics are related to clusters (cell types), where high values are expected.

#### 2 Supplementary Figures

##### 2.1 Simulation study

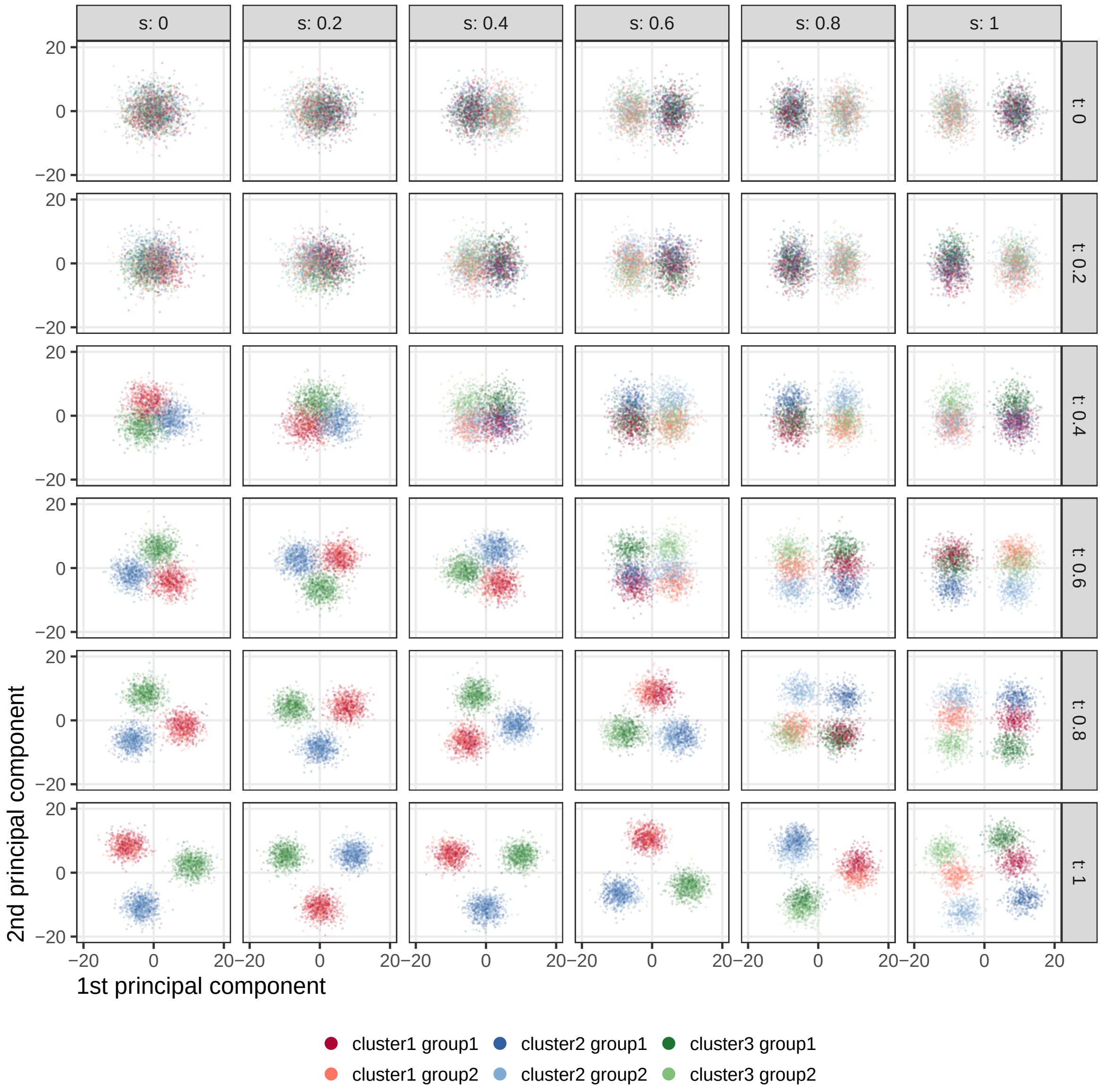

**Fig. S1: PCs across simulations.** Type effect (logFC between groups) and state effect (logFC between conditions) increasing from top to bottom and left to right, respectively; each point represents a cell colored by group and shaded by condition.

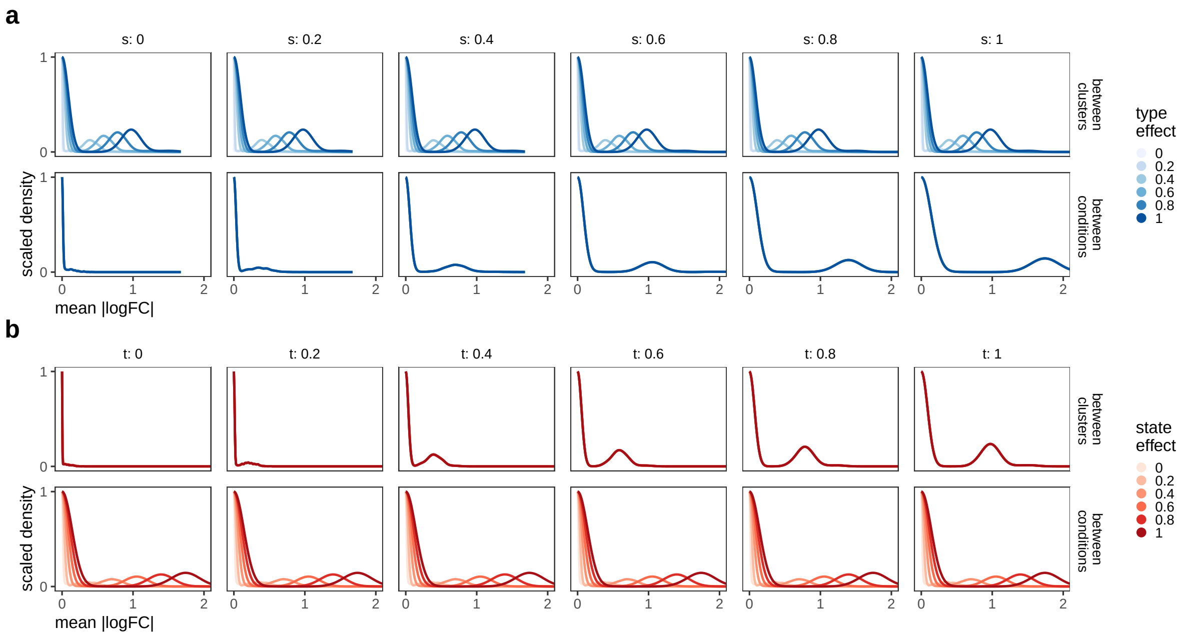

**Fig. S2: Distributions of simulated type and state effects.** Data in both panels are the same, but are stratified/colored by (a) state/type and (b) type/state effect, respectively. Top and bottom row in each panel contain effects between groups and conditions, respectively (see Equation 1). Included are data across 36 simulations with varying effects in $[0, 0.2, 0.4, 0.6, 0.8, 1]$.

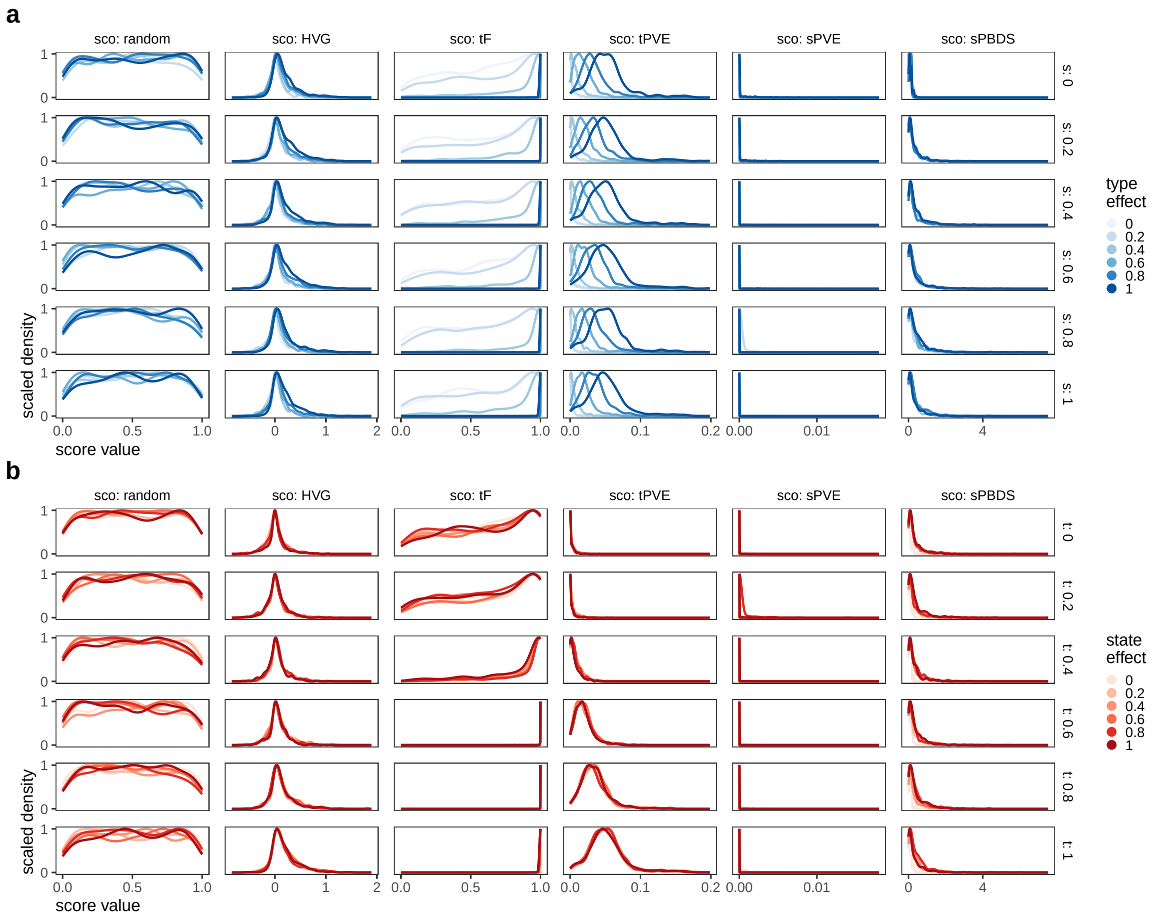

**Fig. S3: Feature score distributions across methods and simulations.** Data in both panels are the same, but are stratified/colored by (a) state/type and (b) type/state effect, respectively. Columns correspond to different feature scoring strategies. Included are data across 36 simulations with varying effects in $[0, 0.2, 0.4, 0.6, 0.8, 1]$, and only genes selected for according to DEnotDS.

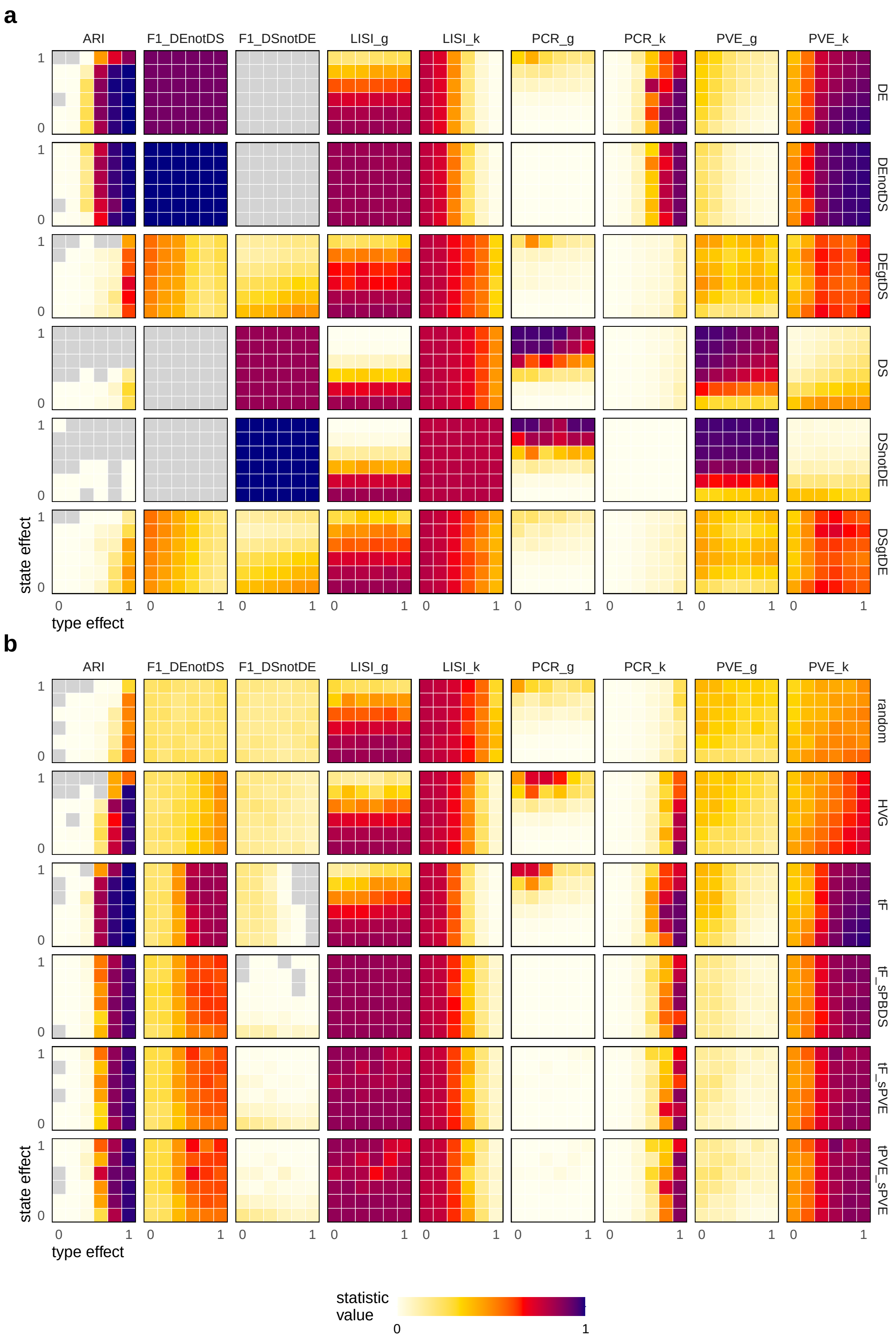

**Fig. S4: Evaluation of feature selections.** Rows = selection methods, columns = evaluation statistics; each panel includes results for a given selection and statistic across 36 simulations with varying type and state effect in $[0, 0.2, 0.4, 0.6, 0.8, 1]$. Results are stratified by (a) simulation parameter-based and (b) other feature selection strategies.

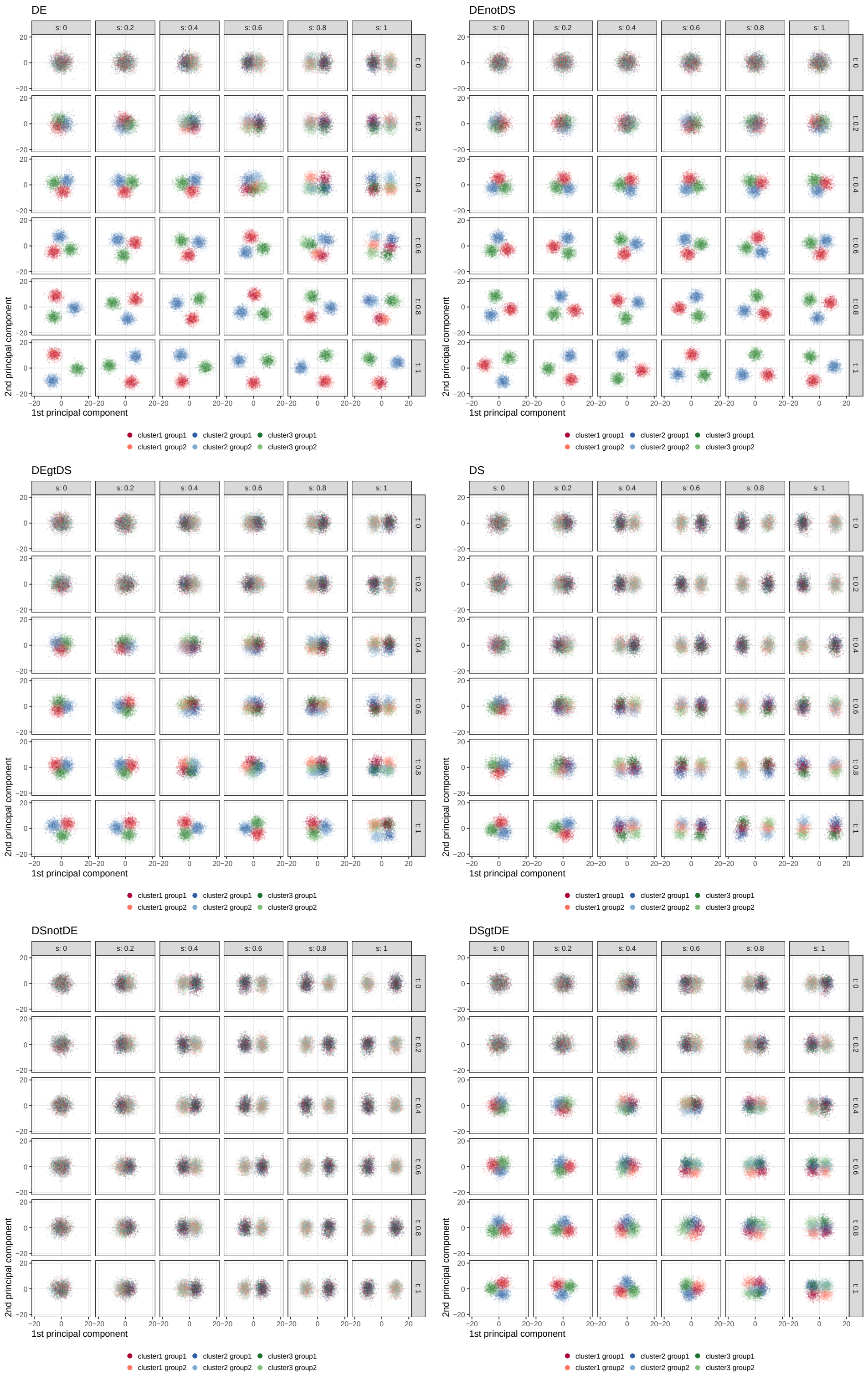

**Fig. S5: PCs based on simulation parameter-based feature selection.** Each panel includes PCs for a given selection across 36 simulations with type and state effect increasing from top to bottom and left to right, respectively; each point represents a cell colored by group (cluster) and shaded by condition.

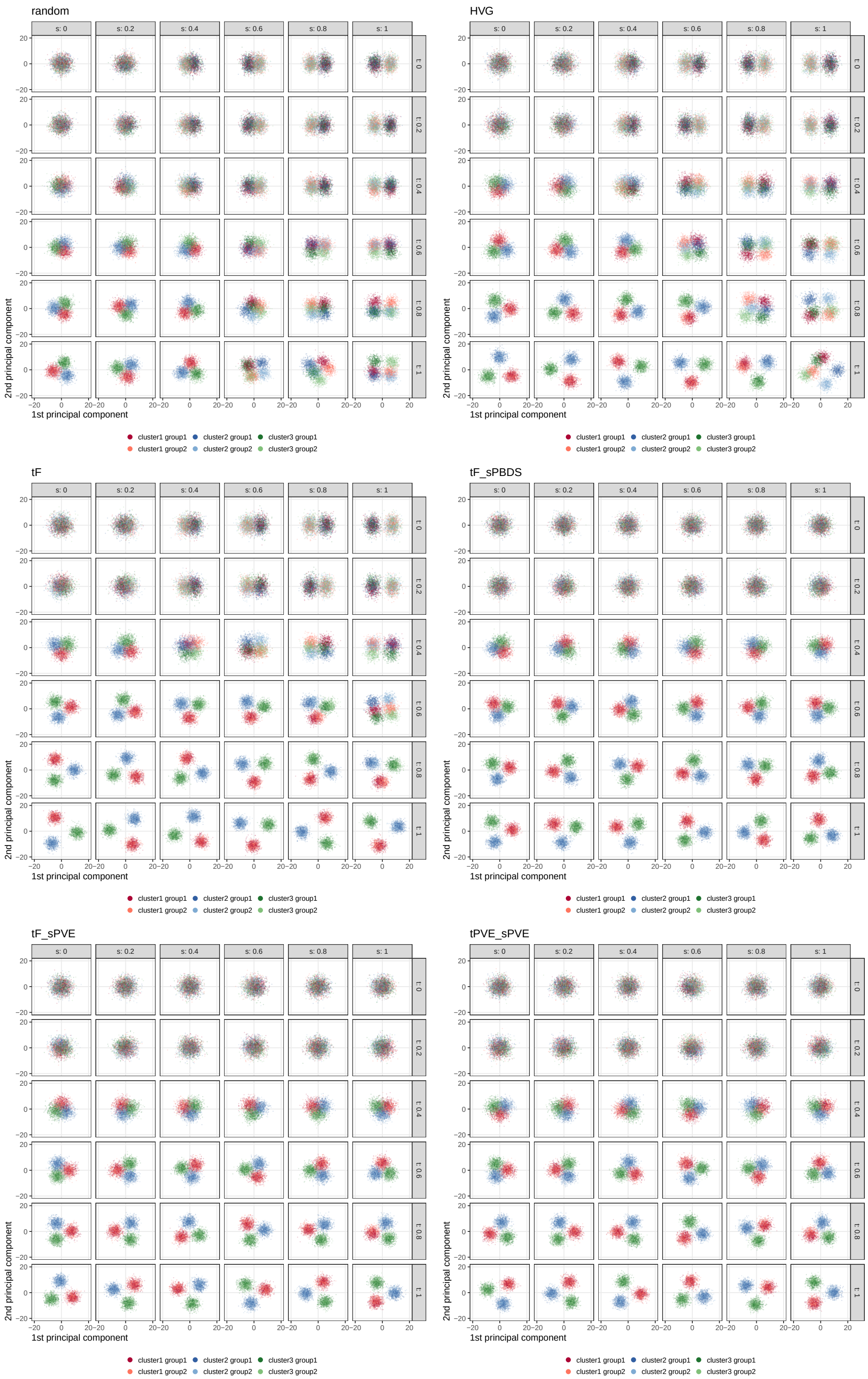

**Fig. S6: PCs based on different feature selection strategies.** Each panel includes PCs for a given selection across 36 simulations with type and state effect increasing from top to bottom and left to right, respectively; each point represents a cell colored by group (cluster) and shaded by condition.

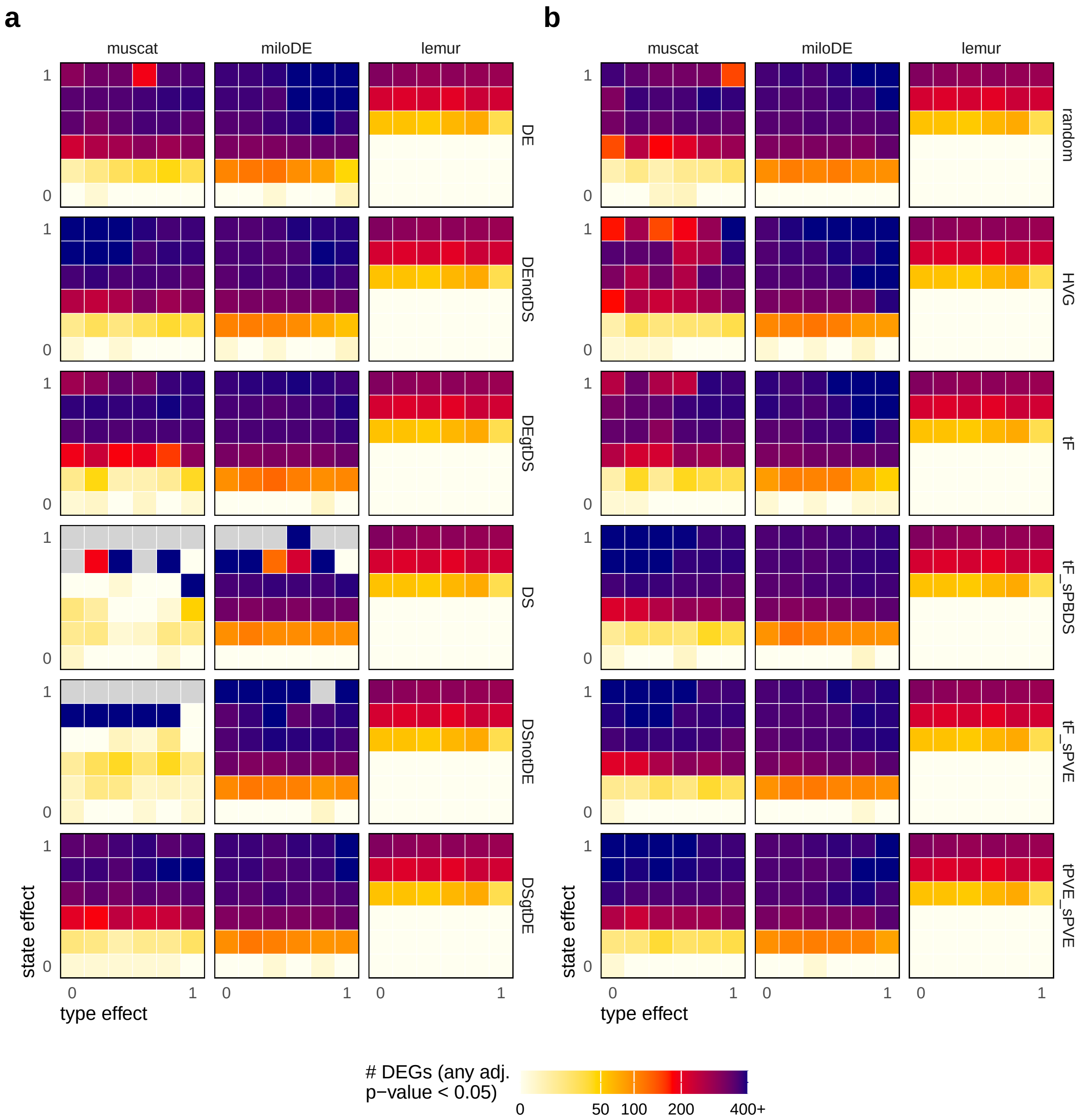

**Fig. S7: Number of genes deemed differentially expressed in any cross-group comparison across simulations, feature selections, and DSA methods.** Rows = selection methods, columns = DSA methods; results are stratified by (a) simulation parameter-based and (b) other feature selection strategies. Heatmap values correspond to the number of genes with an adjusted p-value $<$ 0.05 in any comparison (cluster for muscat, neighborhood for lemur and miloDE). Each panel includes 36 simulations with varying type and state effect in $[0, 0.2, 0.4, 0.6, 0.8, 1]$; heatmap values are square-root scaled for visualization.

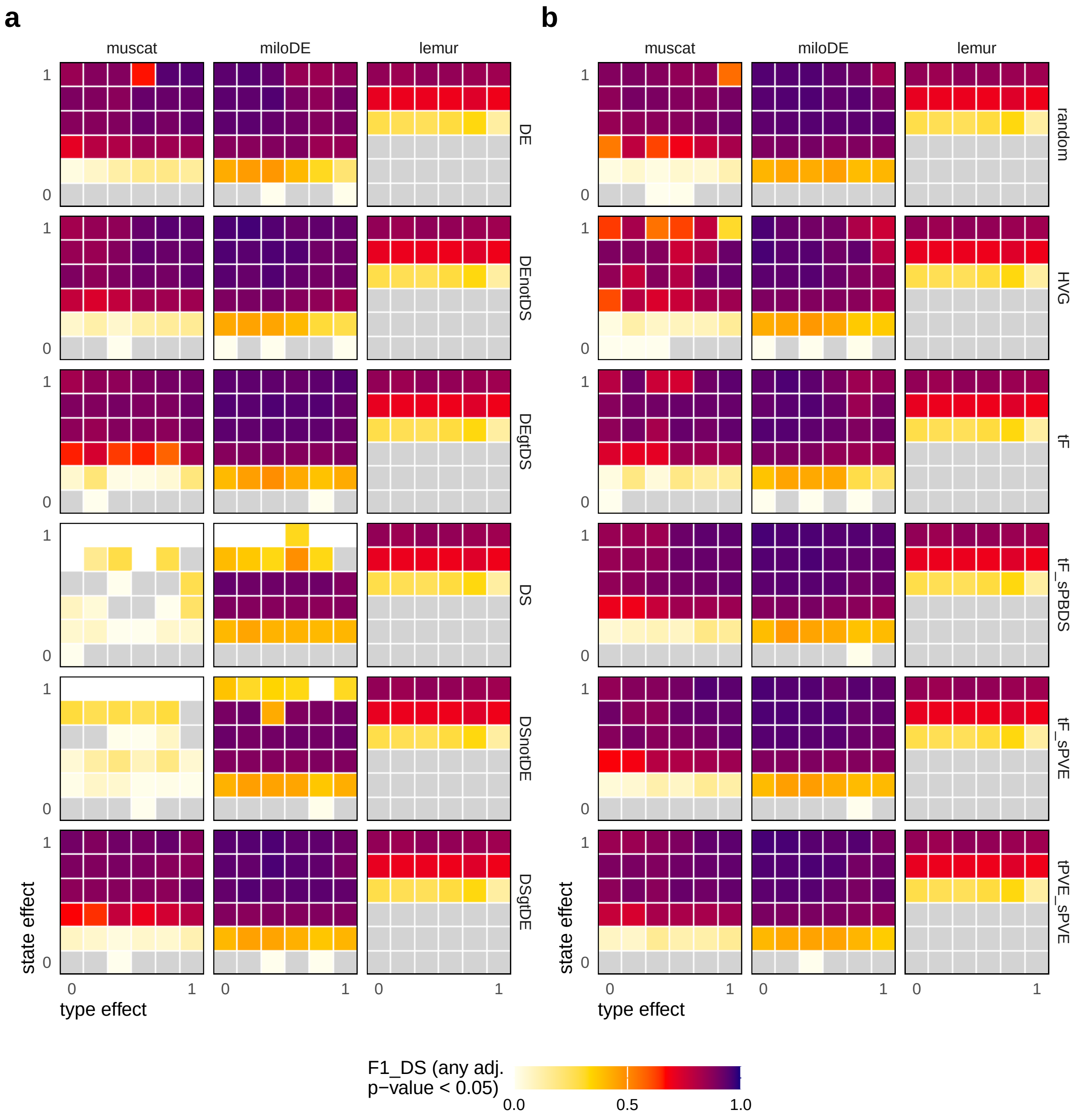

**Fig. S8: Comparison of DSA performance across simulations and feature selections.** Rows = selection methods, columns = DSA methods; results are stratified by (a) simulation parameter-based and (b) other feature selection strategies. Heatmap values correspond to F1 scores (harmonic mean of precision and recall) with DS genes as ground truth ($\text{logFC}_{state} \neq 0$), and genes with an adjusted p-value $< 0.05$ in any cluster (muscat) or neighborhood (lemur, miloDE) as prediction. Each panel includes 36 simulations with varying type and state effect in $[0, 0.2, 0.4, 0.6, 0.8, 1]$.

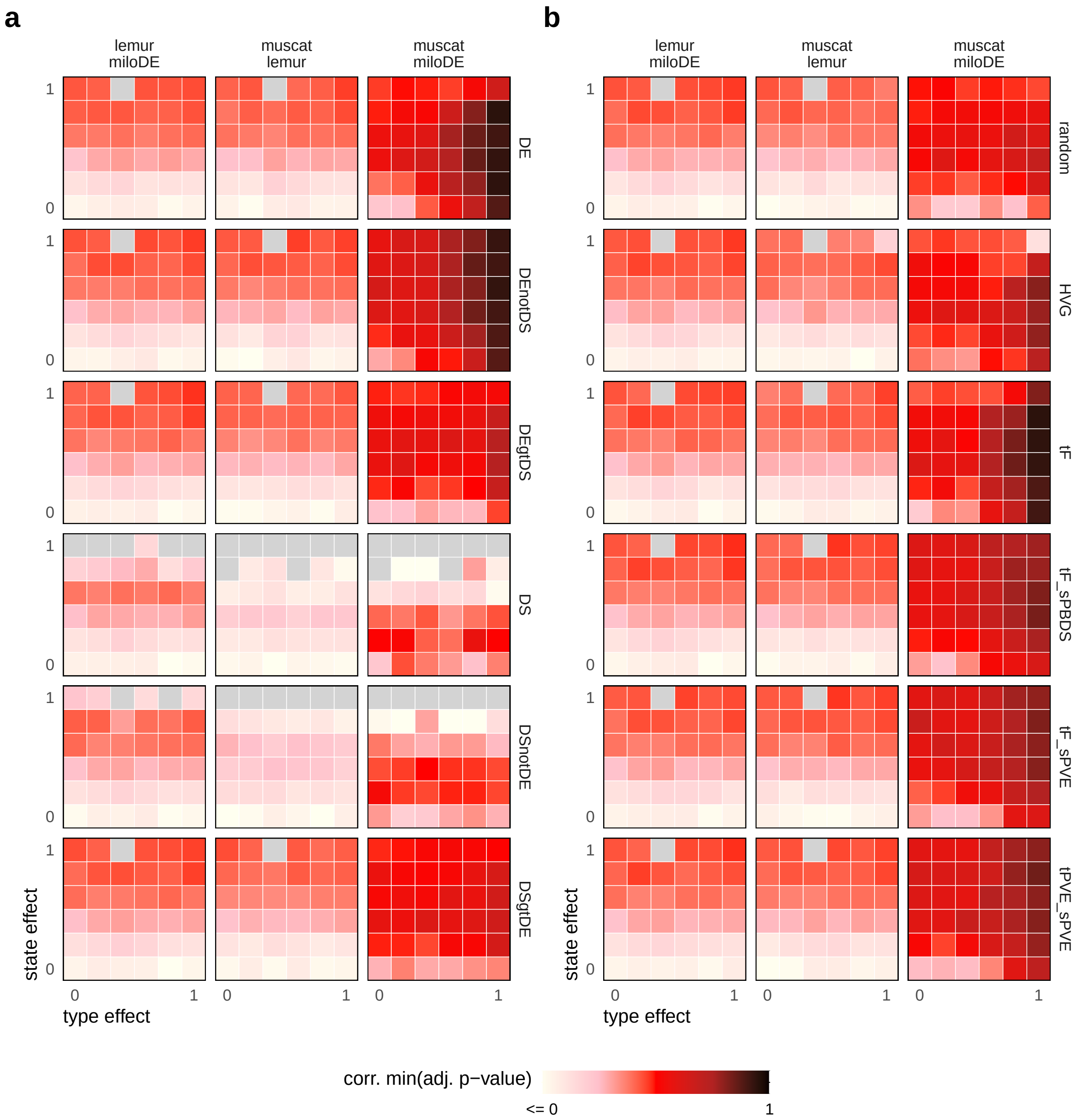

**Fig. S9: Concordance between cluster-based and -free DSA results across simulations and feature selection strategies.** Rows = selection methods, columns = DSA methods; results are stratified by (a) simulation parameter-based and (b) other feature selection strategies. Heatmap values correspond to Spearman correlations between the smallest adjusted p-values across clusters ($\mathtt{muscat}$) or neighborhoods ($\mathtt{lemur}$, $\mathtt{miloDE}$) for each gene. Each panel includes comparisons across 36 simulations with varying type and state effect in $[0, 0.2, 0.4, 0.6, 0.8, 1]$.

#### 2.2 Experimental data

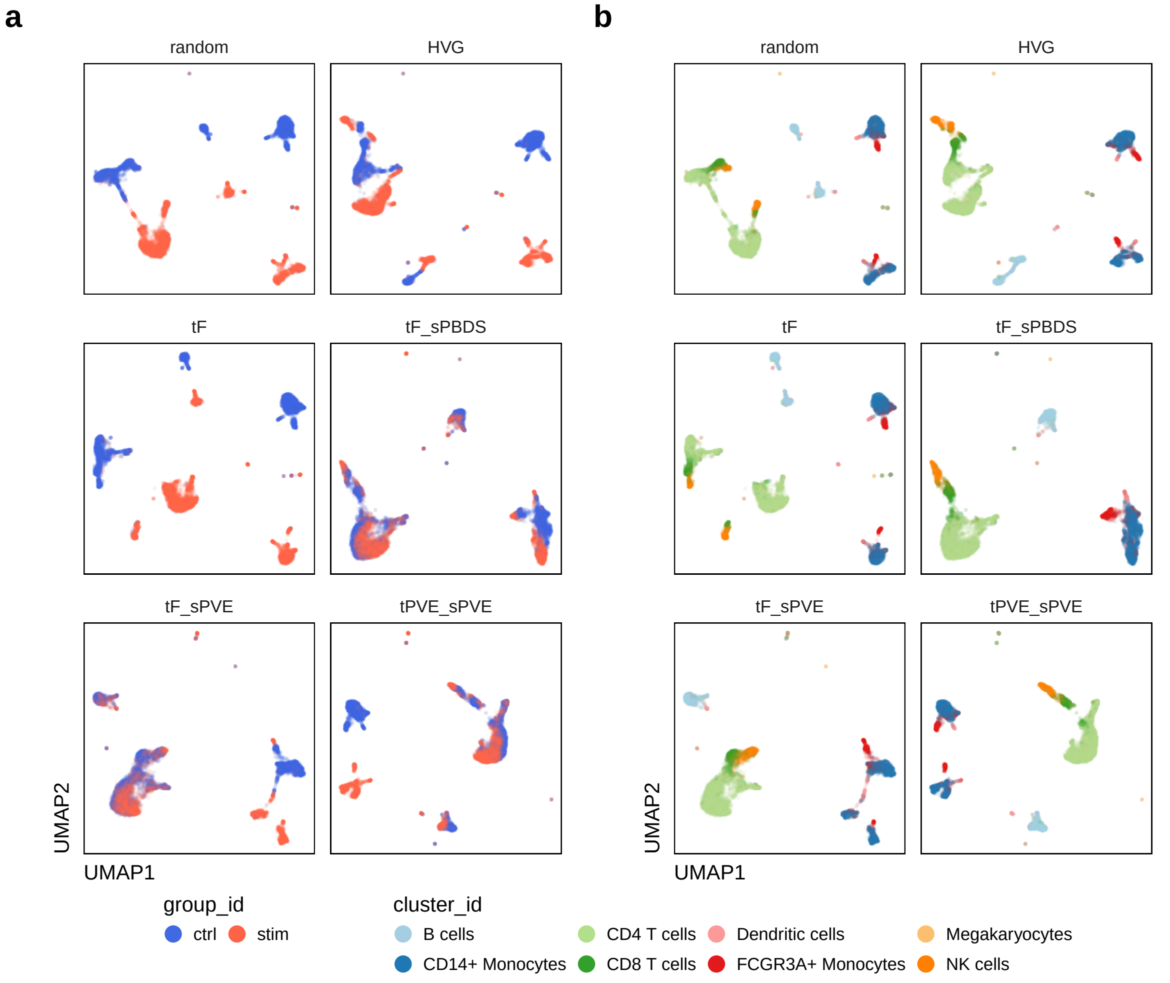

**Fig. S10: UMAP embeddings across feature selection strategies.** Points = cells are colored by condition (a) and subpopulation (b), respectively. Each panel corresponds to the UMAP obtained from PCs based on the 2,000 highest ranked genes for a given selection.

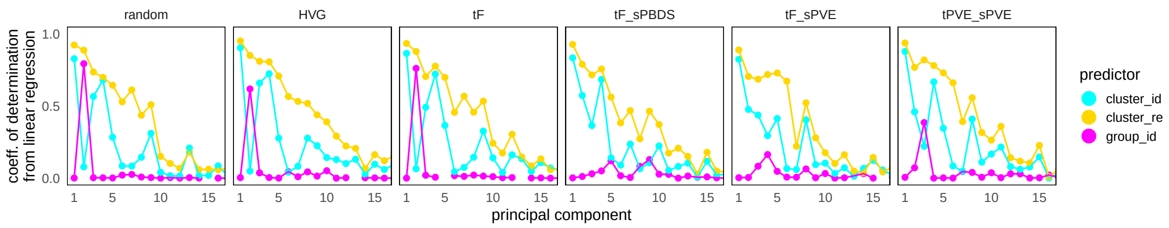

**Fig. S11: Principal component regression against different predictors.** Y-axis values denotes R-squared values from fitting a linear regression model with principal components (PCs) as response, and experimental condition (group_id), true (cluster_id) and inferred (cluster_re) cluster assignments as predictor; different panels are based on different feature selection strategies.

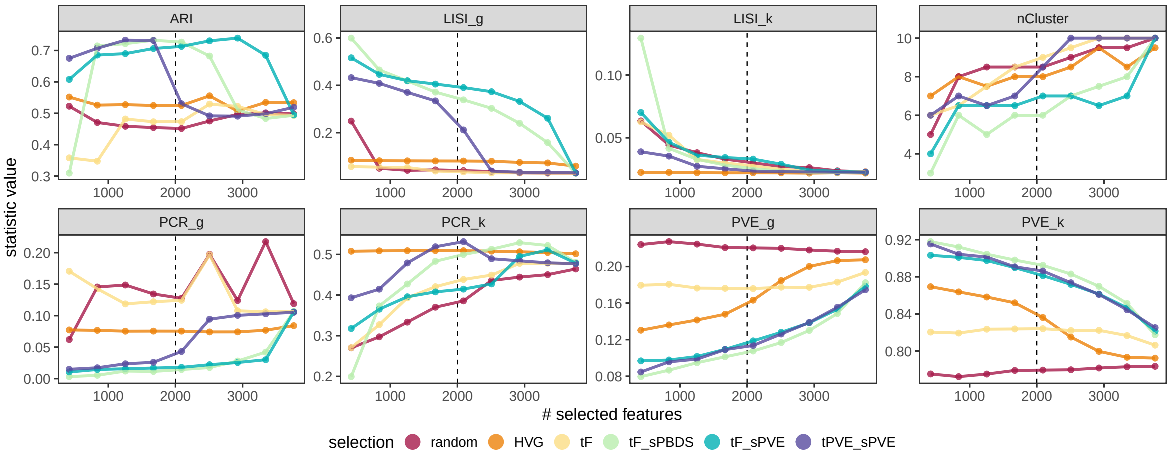

**Fig. S12: Dependence of evaluation statistics on the number of features selected for processing.** Points correspond to data processed based on $10, 20, \ldots, 90\%$ of top-ranked genes for a given selection (= different colors), vertical dashed line at 2,000 corresponds to the number of genes generally selected for.

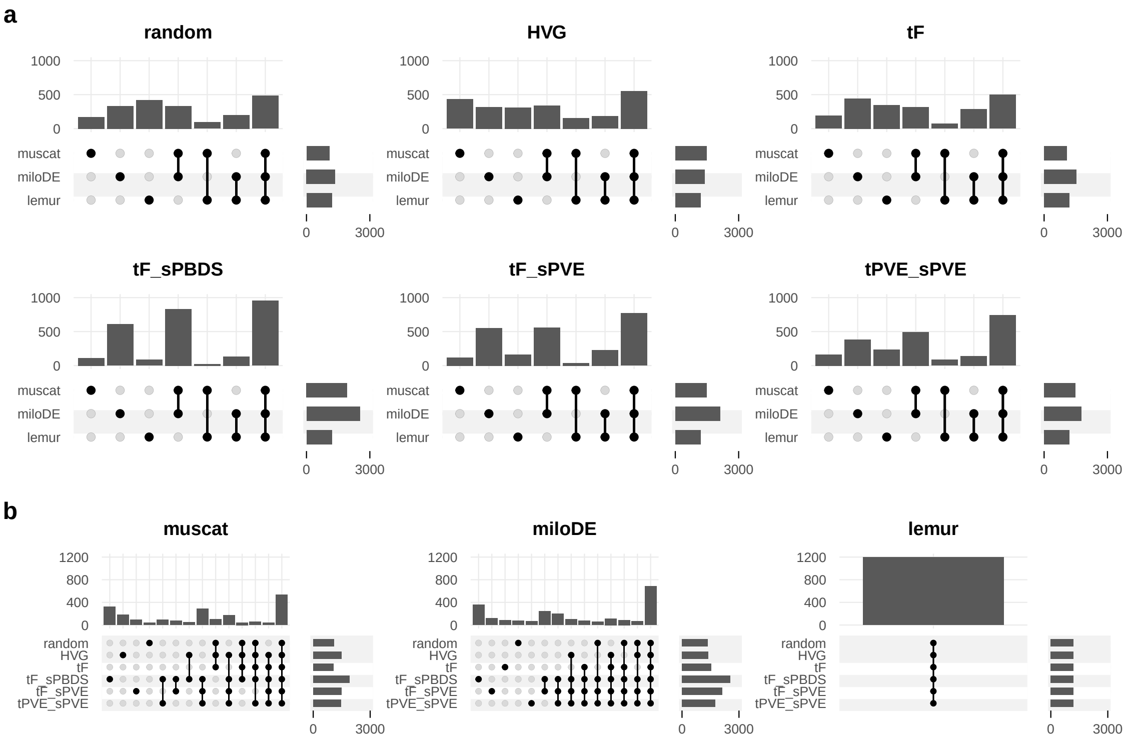

**Fig. S13: Upset plots of DEGs across DSA methods and feature selections.** Genes are deemed differential when their p-value $< 0.05$ in any cluster ($\mathtt{muscat}$) or neighborhood ($\mathtt{lemur}$, $\mathtt{miloDE}$). Both panels show the same data, where in (a) sets = DSA methods, panels = feature selections; and, (b) sets = feature selections, panels = DSA methods.

### 3 Supplementary Methods

#### 3.1 Computation workflow

##### Simulation

*00-get_sim.R* simulates scRNA-seq data using $\mathtt{splatter}$[1], according to input parameters t and s $\in \{0, 0.2, 0.4, 0.6, 0.8, 1\}$, which were passed to $\mathtt{de.facLoc}$ and $\mathtt{cde.facLoc}$ to control gene expression fold changes between clusters (types) and groups (states), respectively. For each set of parameters, we simulated 2,000 genes and $\sim 400$ cells for 6 samples from 2 conditions, each with 3 groups (clusters), resulting in a total of $\sim 2,400$ cells with equal distribution of cells across groups, samples and conditions ($\mathtt{group.prob} = 1/3$ and $\mathtt{condition.prob} = 1/2$).
Simulation parameters were specified via $\mathtt{splatPopSimulate}$ with equal scale factors for groups and conditions ($\mathtt{de.facScale} = \mathtt{cde.facScale} = 0.1$), the probability of genes being differentially expressed between clusters and groups, respectively, both set to 10% ($\mathtt{de.prob} = \mathtt{cde.prob} = 0.1$), a biological coefficient of variation (BCV) underlying common variability across genes of $\mathtt{bcv.common} = 1.5$. A very low sample effect was simulated by setting the scaling factor for the population variance rate parameter to $\mathtt{similarity.scale} = 15$, and one of the eQTL effect parameters to $\mathtt{eqtl.ES.rate} = 30$. To mitigate the dampening of state effects caused by high inter-sample similarity, we amplified the state effect by multiplying $\mathtt{cde.facLoc}$ by 1.2 in each simulation. The seed for random number generation was fixed across all simulations.

##### Preprocessing

*01-pro_dat/sim.R* performs general quality control and preprocessing steps. It keeps features with a count greater than 1 in at least 10 cells in simulated data, and cells with at least 10 detected features. For experimental data, it retains features expressed in at least 20 cells and all cells. The scripts then conducts log-library size normalization using $\mathtt{logNormCounts}$ from $\mathtt{scater}$[2], selects highly variable features (HVGs) using $\mathtt{modelGeneVar}$ from $\mathtt{scran}$[3], while blocking by sample identifier.
Next, principal component analysis (PCA) is performed on HVGs using $\mathtt{runPCA}$ from $\mathtt{scater}$[2], with $\mathtt{ncomponents} = 10$. A shared-nearest neighbor (SNN) graph is built using $\mathtt{buildSNNGraph}$ ($\mathtt{scran}$[3]), with *k=10*. Louvain clustering[4] is then performed with high and low resolution of 2 and 0 for type and state score calculations, respectively, with cluster assignments stored under $\mathtt{colData}$ slot $\mathtt{cluster\_lo/hi}$.

##### Scoring & selection

*02-sco.R* applies a *02-sco_sim/dat-x.R* script to a given simulated/experimental dataset, which computes a variability score for each feature (Table S1), and returns a table of gene-level metadata (including variability scores).
Subsequently, *03-sel.R* calls on a *03-sel_sim/dat-x.R* script, which employs a feature selection strategy (Table S2) – based on computed variability scores, or combinations thereof – to return a logical vector that specifies which features were selected. Here, the top-$n$ ranked features are selected, where $n$ corresponds to the number of ground truth type genes ($\mathtt{DEnotDS}$, i.e., type effect but no state effect) for simulated data, and $n = 2,000$ for experimental data.

##### Reprocessing

*04-rep.R* uses selected features for PCA of a given dataset, and performs SNN graph-based Louvain clustering[4] using $\mathtt{cluster\_louvain}$ ($\mathtt{igraph}$) with $\mathtt{resolution} = 1$. Newly computed cluster assignments are stored under $\mathtt{colData}$ slot $\mathtt{cluster\_re}$. Reprocessed data are then subjected to downstream evaluation and differential discovery.

##### Evaluation

*05-sta.R* applies a *05-sta-x.R* script on reprocessed data to evaluate feature selection and clustering accuracy, as well as overall data structure (Table S3). Depending on the type of metric, and single statistic (e.g., ARI), cluster- and group-level results (e.g., PCR), or gene-level results (e.g., PVE) are returned. For F1 scores, simulated ground truths are considered (c.f. Table S2).

##### DS analysis

*06-das.R* calls on *06-das-x.R* scripts to perform DSA. Here, muscat is applied using cluster assignments from reprocessing (colData slot cluster_re), miloDE uses precomputed PCs, and lemur is run on all data 'as-is'. In any run, *06-das-x.R* also stores unique identifiers (i_nhood) for every instance subjected to cross-group comparison, i.e., clusters (muscat) or neighborhoods (lemur, miloDE); and, computes the number of cells (n_cells) as well as the highest proportion of cells across groups assigned to each instance (p_nhood).

##### Visualization

*08-plt_x-y.R* (*qlt* for experimental data) scripts collect simulation study results of $x$ to generate a plot with identifier $y$. For example, *08-plt_das-n_de.R* plots the number of DEGs from DSA across simulations and feature selections. Similarly, *08-qlt_x-y.R* scripts collect and visualize experimental data results. These are saved to the *plts/sim/* and *plts/dat/* directory, respectively.

##### Miscellaneous

*07-eva.R* is standalone script that is applied to experimental data only. It collects results across all feature selection strategies, select [10, 20, ..., 90%] of top-ranked features, and recomputes evaluation statistics for correspondingly reprocessed data (PCA, clustering).
*09-aes.R* is sources in every R session. It simply fixes the order of feature scores, ground truth-based and other selections, and differential state analysis methods across visualizations.
*10-session_info.R* lists and may be used to install all R packages used (across CRAN, GitHub, and Bioconductor), and write the corresponing sessionInfo output to a *.txt* file.
